# Mixture models reveal multiple positional bias types in RNA-Seq data and lead to accurate transcript concentration estimates

**DOI:** 10.1101/011767

**Authors:** Andreas Tuerk, Gregor Wiktorin, Serhat Güler

## Abstract

Quantification of RNA transcripts with RNA-Seq is inaccurate due to positional fragment bias, which is not represented appropriately by current statistical models of RNA-Seq data. This article introduces the Mix^2^(rd. ”mixquare”) model, which uses a mixture of probability distributions to model the transcript specific positional fragment bias. The parameters of the Mix^2^ model can be efficiently trained with the Expectation Maximization (EM) algorithm resulting in simultaneous estimates of the transcript abundances and transcript specific positional biases. Experiments are conducted on synthetic data and the Universal Human Reference (UHR) and Brain (HBR) sample from the Microarray quality control (MAQC) data set. Comparing the correlation between qPCR and FPKM values to state-of-the-art methods Cufflinks and PennSeq we obtain an increase in R^2^ value from 0.44 to 0.6 and from 0.34 to 0.54. In the detection of differential expression between UHR and HBR the true positive rate increases from 0.44 to 0.71 at a false positive rate of 0.1. Finally, the Mix^2^ model is used to investigate biases present in the MAQC data. This reveals 5 dominant biases which deviate from the common assumption of a uniform fragment distribution. The Mix^2^ software is available at http://www.lexogen.com/fileadmin/uploads/bioinfo/mix2model.tgz.

## 1 Introduction

RNA-Seq has established itself as a popular alternative to microarrays for the quantification of RNA transcripts. In contrast to microarrays, which measure the quantity of an RNA transcript by hybridization to a transcript specific oligonucleotide, RNA-Seq generates cDNA for fragments of the RNA transcript, which are sequenced by a next generation (NGS) sequencer. One advantage of RNA-Seq over microarrays is that it does not require prior knowledge of the nucleotide sequence of the RNA transcript, which is needed to produce a transcript specific hybridization probe, and that it can therefore detect and quantify novel RNA transcripts. In addition, quantification by RNA-Seq covers a wider dynamic range since microarrays suffer from signal saturation resulting in the truncation of abundance estimates for highly abundant transcripts [3].

Despite these advantages, obtaining accurate transcript quantification measurements from RNA-Seq has proven difficult. One of the main reasons for the inaccuracy is the failure of the statistical models used in the derivation of the measurements to properly represent biases inherent in RNA-Seq data. The statistical model of Cufflinks [17], for instance, assumes that the cDNA fragments generated by RNA-Seq are uniformly distributed along the transcripts. In reality, however, this assumption is rarely fulfilled and quantification measurements by Cufflinks are therefore often inaccurate.

One type of bias affecting transcript quantification from RNA-Seq data is the result of a preference of the fragmentation, i.e. the process that generates cDNA fragments from RNA transcripts, to produce fragments at certain positions within the transcript, e.g. at the start and/or at the end of the transcript. Hence, this type of bias is referred to as positional bias [14]. Positional bias can also be caused by a bias in the RNA itself, for instance, due to RNA degradation which results in a shortening of the RNA. Another kind of bias in RNA-Seq is introduced during ligation, amplification and NGS sequencing [2]. This bias is correlated to the RNA sequence of a transcript and is therefore termed sequence specific bias [14]. The present article focuses on the first type of bias, i.e. the positional bias, and develops a model, the Mix^2^ model (rd. ”mixquare”), which performs significantly better than Cufflinks [17] and PennSeq [4] both on synthetic data and the Universal Human Reference (UHR) and brain (HBR) samples from the Microarray Quality Control (MAQC) experiment [10].

The inclusion of bias models into the statistical models of RNA-Seq data has been investigated before. In [8] a model is proposed to account for the variability in read counts depending on the sequence surrounding the start of a fragment. The intention is similar to that of the fragment specific bias model in [14], which also investigates the sequences surrounding the start of a fragment. Similar to Cufflinks [17], the generative models in [6] and [5] describe the probability of observing a fragment for a given transcript. In addition to Cufflinks [17], however, [6] and [5] introduce additional hidden variables and use a bias model, which is derived from the global bias observed in the complete RNA-Seq data set. The method described in [21], on the other hand, is a model for gene read counts, which models bias by exon specific weights, which are estimated both for the complete data set and for individual genes. In [19] the authors focus on RNA-Seq data with 5’ bias which is the result of RNA degradation and use an exponential model for the fragment distributions. The model proposed in [9] is, again, a model for the read counts of a gene. Here the read counts are modelled by a quasi-multinomial distribution with a parameter that can be adapted to account for over and under dispersion. The authors in [14] propose a model both for sequence specific and positional bias. For the positional bias a non-parametric model is used, which can theoretically be trained with the EM algorithm. However, due to the large number of variables, this is feasible only for a simplified version of the model. In practice, biases models in [14] are estimated for few transcript length classes, for which statistics are collected in a small number of bins. The model of [14] has been implemented as an optional extension of Cufflinks [17] and will henceforth be referred to as Cufflinks with bias correction. The model developed in PennSeq [4] is again non-parametric and the large number of variables makes its training computationally prohibitive. For this reason, the bias model of PennSeq [4] is not included in the parameter update but is approximated by the overall bias in a gene locus and by the transcript specific reads.

Similar to some of the other methods mentioned above, the Mix^2^ model is a generative model of the probability of a fragment in an RNA-Seq data set. In contrast to the positional bias model in Cufflinks [14] and PennSeq [4], however, the Mix^2^ model is parametric, which considerably simplifies adaptation of the Mix^2^ parameters. On the other hand, due to the representation of the transcript specific fragment bias by mixtures of probability distributions, the Mix^2^ model is flexible enough to allow for multiple positional biases of arbitrary complexity.

## 2 Materials and methods

### 2.1 The Mix^2^ model: a mixture of mixtures

An essential part of next generation sequencing (NGS) is the library preparation. This process takes an RNA sample and produces a library of short cDNA fragments, each corresponding to a section of an RNA transcript. The cDNA fragments are sequenced by an NGS sequencer resulting in single or paired end reads which are mapped to a reference genome. Hence, the probability *p*(*r*) of a fragment *r* can be interpreted as the probability of its genomic coordinates. In a genomic locus the probability *p*(*r*) is the superposition of the fragment distributions *p*(*r*|*t* = *i*) for the *N* transcripts in the locus, i.e.

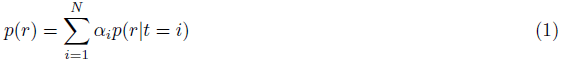

where *α*_*i*_ is the relative abundance of transcript *t* = *i*, i.e. the probability that transcript *t* = *i* generates any fragment, and *p*(*r*|*t* = *i*) is the probability that transcript *t*= *i* generates fragment *r*. An estimate for the concentration of transcript *t* = *i* is obtained by normalizing the relative abundance *α*_*i*_, yielding the RPKM [12] or FPKM values [17].

The Mix^2^ model uses a mixture model for *p*(*r*|*t* = *i*), i.e.

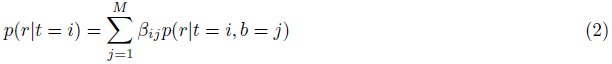

where *M* is the number of mixture components and the hidden variable *b* = *j* represents a ”building block” of the fragment distribution *p*(*r*|*t* = *i*). In addition, for each *i*, the β_*ij*_ sum to one. Since *p(r)* is itself a mixture of the *p*(*r*|*t* = *i*) with weights *α*_*i*_ this implies that *p*(*r*) is a mixture of mixtures, which explains the name of the Mix^2^ model.

In the following *p*(*r*|*t*=*i*) is factorized according to

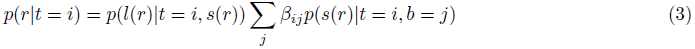

where *s*(*r*) and *l*(*r*) are the start and length of fragment *r* and *p*(*s*(*r*)|*t* = *i*, *b* = *j*) are Gaussians whose means *μ*_*ij*_ are placed equidistantly along the transcript. In transcript coordinates the means *μ*_*ij*_ are given by

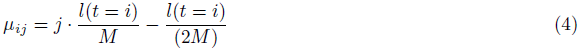

and the standard deviations are, independent of *j*, set to

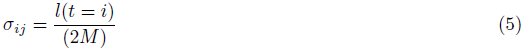

where *l*(*t* = *i*) is the length of transcript *t* = *i*. The Gaussians are, furthermore, normalized such that their sum over the possible fragment starts *s* = 1,...,*l*(*t*) equals one. An example for such a Mix^2^ model can be found in Figure 1. Figure 1 (a) shows the mixture weights *β*_*ij*_ whereas Figure 1 (b) shows the weighted Gaussians, *β*_*ij*_*p*(*s*|*t* = *i*, *b* = *j*), and the sum of the weighted Gaussians, *p*(*s*|*t* = *i*). The distributions in Figure 1 (b) are given in transcript coordinates for a transcript of 2000 bps length, while the longer dashed curve in Figure 1 (c) shows *p*(*s*|*t* = *i*) in genome coordinates. The locus in Figure 1 (c) contains two transcripts which share a common junction. The shorter of the transcripts in Figure 1 (c) has the same set of β_*ij*_ as in Figure 1 (a) but is only 1000 bps long. The relative abundances of the long and short transcript are 0.7 and 0.3, respectively, which results in the overall distribution p(s) given by the solid curve in Figure 1 (c). In comparison, Cufflinks [17] can, for this locus, only model fragment start distributions *p*(*s*|*t* = *i*) as visualized by the dashed curves in Figure 1 (d) and is therefore inappropriate for 5’ biases as the one in Figure 1 (c).

**Figure 1:**
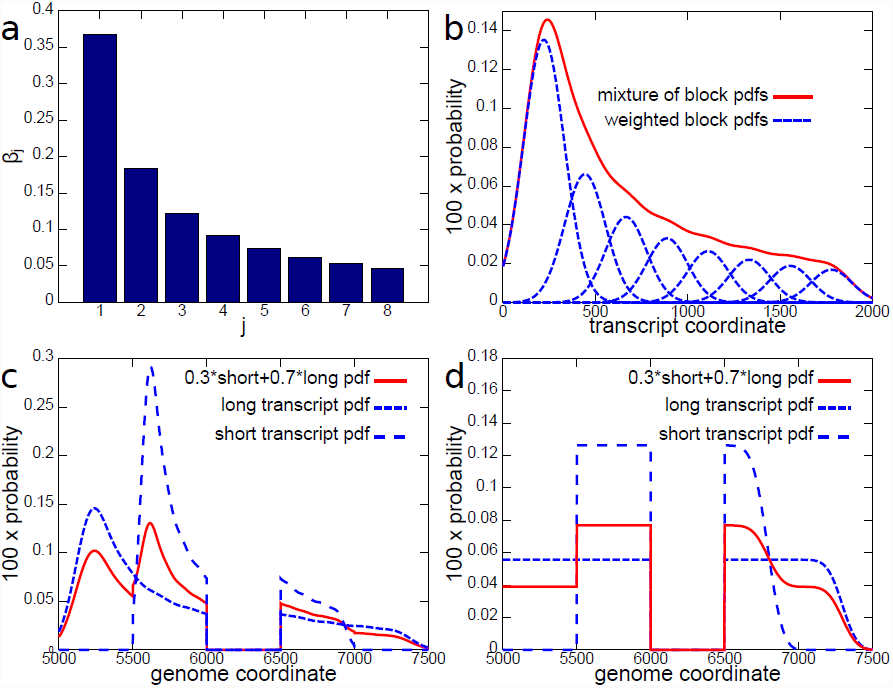
Fragment start distributions modelled by Mix^2^ model and Cufflinks. (a) Set of eight *ß*_*ij*_. (b) Dashed curves: *ß*_*ij*_*p*(*r*|*t* = *i*, *b* = *j*). The *p*(*r*|*t* = *i*, *b* = *j*) are Gaussians equidistantly distributed along a 2000 bps transcript. Solid curve: *p*(*r*|*t* = *i*). All in transcript coordinates. (c), (d) Fragment distributions in locus with two transcripts, 1000 bps and 2000 bps long, sharing one junction. Long and short transcripts start 5000 bps and 5500 bps from beginning of locus contig. Junction starts at 6000 bps, extends to 6499 bps. Dashed curves: *p*(*r*|*t* = *i*) for Mix^2^ model (c), Cufflinks (d), long and short transcript. Solid curve: *p*(*r*) for Mix^2^ model (c), Cufflinks (d). All in genome coordinates.

### 2.2 Parameter tying

The mixture weights β_*ij*_ determine the shape of the fragment distribution of transcript *t*=*i*. Thus, if some transcripts have a similar distribution then their *β*_*ij*_ can be tied between the transcripts. As will be shown in the experiments in Section 3.2, this can lead to increased accuracy in the abundance estimates. Consider, for instance, the fragment start distribution of the Cufflinks model in Figure 2(a). Here the distributions are very similar for transcripts with 2000 bps and 3000 bps length and for transcripts with 700 bps and 1000 bps length. In this situation therefore, for the Mix^2^ model, these four transcripts could be separated into two groups where the transcripts within each group share the same mixture weights *β*_*ij*_. In general this leads to the scenario, where each transcript *t*= *i* has an associated group *g* = *k* and the distributions *p*(*r*|*t* = *i*) of transcripts within this group share the same β_*ij*_.

**Figure 2:**
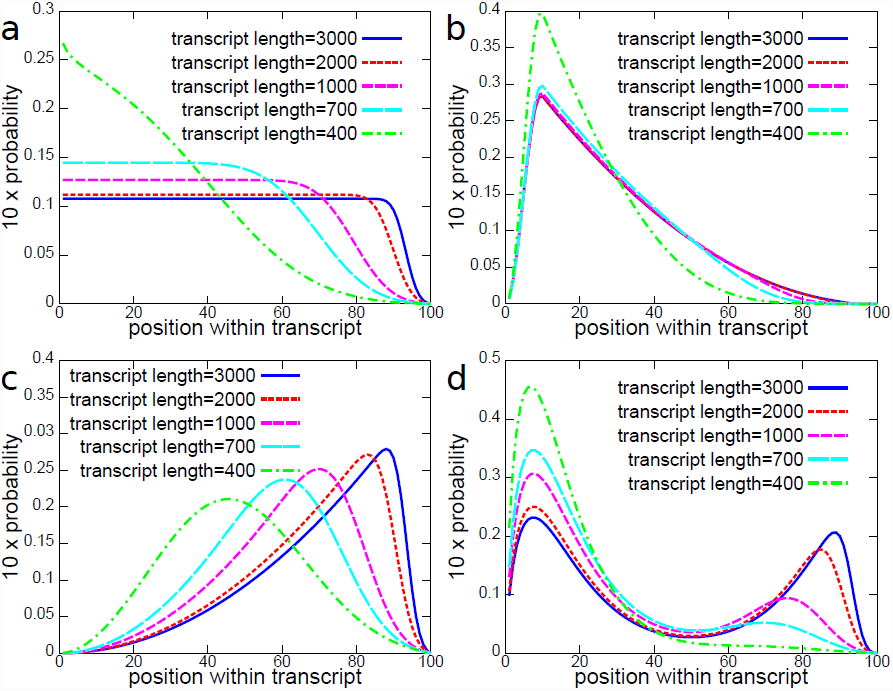
Transcript length dependent fragment start distributions in artificial data. X-axis is position within transcript in percent. Distributions are derived from an initial distribution, stretched to transcript length. Subsequent multiplication with fragment start distribution of Cufflinks (a) and renormalization. (a) Distributions for uniform initial distribution. Corresponds to Cufflinks model for Gaussian fragment length distribution with mean 200 and standard deviation 80. Bias in (a) is referred to as Cufflinks bias. Distributions derived from initial distribution with 5’ bias (b), 3’ bias (c) and 5’+3’ bias (d).

Multiple factors might influence the similarity of fragment start distributions. Here, only gene membership and transcript length are investigated. The rationale for choosing these two properties is that even if fragments are uniformly distributed along a transcript immediately after fragmentation, fragment size selection introduces the transcript length dependent bias in Figure 2 (a). On the other hand, transcripts belonging to the same gene can share a substantial part of their sequence and therefore potentially exhibit similar fragmentation properties.

Based on these two criteria, in the experiments in Section 3.1 transcripts within genes are manually separated into several groups. For larger number of genes, however, this is impractical. Therefore in Section 3.2 a clustering scheme was employed which automatically separates the transcripts within a gene into different groups. In this scheme, transcripts were initially placed into groups according to their length where the 7 transcript length boundaries were equidistantly distributed on the log scale between 300 and 5000. Subsequently, groups were merged until each group had at least 20 valid reads and there was at most one group containing a single transcript. Groups were merged according to their distance, which was calculated as follows

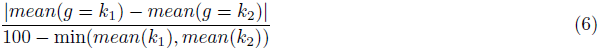

where *mean*(*g* = *k*) is the average length of transcripts in group *g* = *k*. The two closest groups were merged first. This type of tying is referred to as group tying. If all transcripts within a gene are placed into a single group this is referred to as global tying. Hence, Figure 1 (c) shows an example for a Mix^2^ model with global tying, because all the transcripts in this locus share the same set of weights *β*_*ij*_.

### 2.3 Parameter estimation

The relative abundances *α*_*i*_ in the Mix^2^ model can be update with the EM algorithm in the usual manner, as implemented, for instance, in Cufflinks [17]. This update formula is given in Section 1.1 of the supplement. For the transcripts in group *g* = *k* the *β*_*ij*_ = *β*_*kj*_ can be updated with the EM algorithm as follows

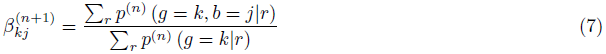

where

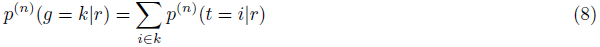

is the sum over the transcripts *t* = *i* in group *g* = *k* and a similar equation holds for *p*^(*n*)^(*g* = *k*,*b* = *j* |*r*). Here 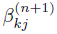 and *p*^(*n*)^ (•) are the mixture components and posterior probabilities after the *n* + 1-th and after the *n*-th iteration, respectively. Iterations are repeated until changes in the model parameters fall below a predefined threshold. An example of the convergence of the Mix^2^ model parameters can be found in Section 1.2 of the supplement. From (7) the EM update formulas for a Mix^2^ model without parameter tying and with global tying are readily deduced. This derivation together with the derivation of (7) can be found in Section 1.1 of the supplement.

The previous discussion assumes that a fragment *r* maps uniquely to the reference. If, on the other hand, fragment *r* has multiple hits *H*(*r*) on the reference, then the following holds

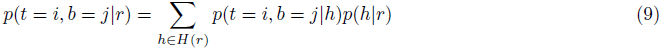

and the update formula for *ß*_*kj*_, equation (7), becomes

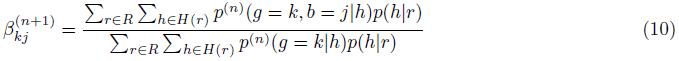

Typically, *p*(*h*|*r*) is set to 1/#*H*(*r*) [12] but can, in principle, be estimated from the data. However, in the experiments on the MAQC data in Section 3.2 no distinction is made between single-and multi-mapping fragments and each hit of a multi-mapping fragment is treated as an individual fragment. Even under this simplifying assumption does the Mix^2^ model yield significantly better results than the compared methods.

## 3 Results

The experiments in this section compare the Mix^2^ model to Cufflinks 2.2.0 with bias correction [14], without bias correction [17] and to PennSeq [4]. Cufflinks and PennSeq were chosen in these experiments because Cufflinks is one of the most widely used methods for the estimation of transcript abundances, while PennSeq has recently been shown to outperform a large number of publicly available abundance estimation methods, including the bias models proposed in [9], [19], [13] and [11].

In the following, the means of the probability distributions *p*(*s*|*t* =; *i*, *b* = *j*) where placed equidistantly along the transcript as in (4) and the standard deviations were set as in (5). The mixture weights and abundances were initialized uniformly, i.e. *β*_*ij*_ = 1/*M* and *α*_*i*_ = *1/N*, where *M* is the number of mixtures in the *p*(*r*|*t* = *i*) and *N* is the number of transcripts in the gene.

### 3.1 Experiments on artificial data

One of the main intentions of the experiments in this section was to find the optimal number of mixtures for the Mix^2^ model depending on the data and the type of model. For this purpose, the 4 transcript length dependent fragment biases in Figure 2 were studied on 7 test genes from GRCh37 Ensembl annotation v75. The genes were selected to represent abundance estimation problems of varying complexity and contain between 4 and 15 transcripts as well as the main variants of differential splicing. Names and properties of these genes are summarized in Table 1 in the supplement. For each of the 7 genes, 200 sets of abundances (*α*_1_,...,*α*_*N*_) were sampled uniformly, according to the Dirichlet distribution. Subsequently, for each of the 200 sets of abundances 500, 1000, 5000 and 10000 fragments were sampled from the superposition (1), where the *p*(*r*|*t* = *i*) belong to one of the 4 bias models in Figure 2. These biases are referred to as Cufflinks bias (a), 5’ bias (b), 3’ bias (c) and 5’+3’ bias (d), where the Cufflinks bias is the fragment start distribution of the Cufflinks model for a fragment length distribution with mean 200 bps and standard deviation 80 bps. The biases in Figure 2 are derived by stretching an initial distribution to the length of the transcript. Subsequently the stretched distribution is multiplied by the Cufflinks bias of the transcript length and renormalized. This explains why the 5’ tails of the biases in Figure 2 become increasingly heavy for shorter transcripts. In the supplement, Section 2 contains a brief discussion of how the Cufflinks bias is derived from the Cufflinks model and Figures 1, 2, 3 and 4 show examples for the coverage resulting from sampling the biases in Figure 2. Finally, the fragment lengths *l*(*r*) were sampled from a Gaussian with mean 200 bps and standard deviation 80 bps and the resulting fragments were then converted into 50bps paired-end reads and written to a SAM file [7]. Thus, 800 data sets were generated per gene and sample size or equivalently 1400 data sets per bias model and sample size resulting in a total of 22400 data sets. On each of these data sets the Mix^2^ model was run without tying as well as with group and global tying, where the number of mixtures ranged from 2 to 6 and included 8, 10 and 20. Hence a total of 537600 experiments were performed with the Mix^2^ model on the artificial data. Thus, while the number of genes in these experiments might seem small the combinatorial explosion of the possible parameter combinations leads to a computational task of sizable proportions.

The difference between true and estimated relative abundances is measured with the L_1_ distance in this section, which is the sum of the absolute differences between true and estimated abundances and can therefore be interpreted as the accumulated error over the complete gene. The L_1_ distance of two probability distributions lies between 0 and 2.

The fragment length distribution was assumed to be given for Cufflinks and the Mix^2^ model and was therefore set to a Gaussian with mean 200 bps and standard deviation 80 bps. It would have been possible to estimate the fragment length distribution from the test data but the difference between true and estimated distribution is minor.

For the Mix^2^ model with group tying the transcripts of a gene were placed into one of two different groups separating long from short transcripts. The groups were chosen ad hoc by visual inspection of the gene annotation independently of the fragment bias. This ad hoc approach is replaced by the clustering procedure from Section 2.2 in the experiments in Section 3.2.

First, the experiments in this section addressed the question whether knowledge about the data is necessary in order to choose an appropriate number of mixtures *M* for the Mix^2^ model. In particular, we wanted to know whether it is necessary to obtain information about the bias, gene or sample size prior to optimization or whether the number of mixtures can be optimized independent of these data properties. We restricted ourselves to optimization on the gene level as brute-force optimization on the transcript level would have significantly increased the number of experiments. Figure 3(a) shows the dependence of the L_1_ distance averaged over all combinations of bias, gene and sample size on the level of optimization. Thus, the three left-most bars in Figure 3(a) show the average L_1_ distance when choosing an optimal number of mixtures for each combination of bias, gene and sample size. The three right-most bars, on the other hand, show the average L_1_ distance after a single optimal number of mixtures has been chosen for all combinations of bias, gene and sample size. Figure 3(a) therefore implies that there is little difference between the average L_1_ distance obtained for different optimization levels and that the number of mixtures can therefore be optimized without referring to gene, sample size or bias. In the studied range the sample size, in particular, appears to play a minor role in the optimization as the average L_1_ distances of the optimization at gene/sample size level and gene level is almost identical. This can also be seen from Figure 3(b) which shows the dependence of the L_1_ distance on the sample size and the number of mixtures. It can, furthermore, be seen that the minimal average L_1_ distance for a Mix^2^ model without tying is obtained for 3 mixtures, while for a Mix^2^ model with group and global tying the minimum is obtained with a number of mixtures between 4 and 10. These optimal numbers of mixtures are independent of the sample size. Figure 3(b) shows furthermore that the effect of tying increases with the sample size. While the difference between the average L_1_ distances of a Mix^2^ model with global and group tying is relatively moderate for sample sizes of 500 and 1000 it becomes more pronounced for samples sizes of 5000 and 10000.

**Figure 3:**
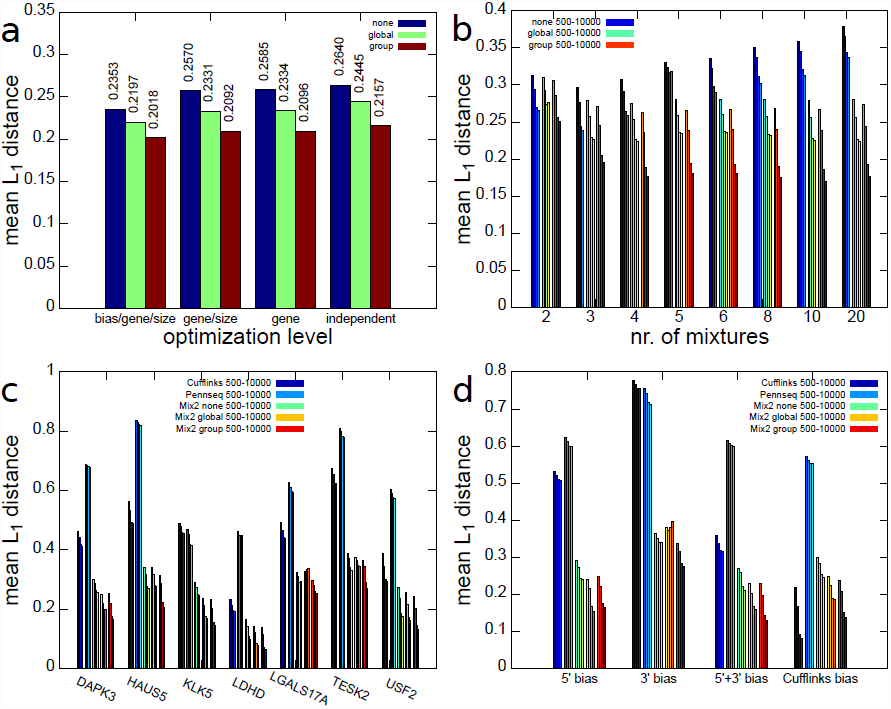
Average L_1_ distance on artificial data between true and estimated abundances for Cufflinks, PennSeq and Mix^2^ model. (a) Dependence of L_1_ distance on level of optimization of mixture number. Each group of 3 blocks gives the average L_1_ distance for Mix^2^ model without tying, with global and with group tying. (b) Dependence of L_1_ distance on number of mixtures and sample size. First four bars for each mixture number are Mix^2^ model without tying, four bars in the middle and on the right Mix^2^ model with global and group tying. Bars within each group correspond to 500, 1000, 5000 and 10000 fragments. L_1_ distance for Cufflinks, PennSeq and Mix^2^ model on 7 test genes (c) and 4 biases (d). First and second 4 bars are Cufflinks and PennSeq.

Figures 3(c) and (d) present a comparison between Cufflinks, PennSeq and the Mix^2^ model, where the latter uses 4 mixtures for global and group tying and 3 mixtures without any tying. Figure 3(c) shows that there is a certain variability between the accuracy obtained for the different genes. Interestingly, the gene with the highest number of transcripts, which is USF2 with 15 transcripts, is not the most difficult gene. Instead HAUS5 and TESK2 with 10 and 7 transcripts yield higher L_1_ distances for all methods. With the exception of KLK5, PennSeq appears to perform worse than Cufflinks on the artificial data. In comparison to Cufflinks and PennSeq, the Mix^2^ model with and without tying performs considerably better. This higher accuracy of the Mix^2^ model is also demonstrated in Figure 3(d) which shows the dependence of the average L_1_ distance on the type of bias. The Mix^2^ model is more accurate than Cufflinks for all bias types apart from the Cufflinks bias. The latter, however, has to be expected as data with Cufflinks bias fit the model in Cufflinks perfectly. For all bias types except the 5’ bias, group tying can be seen to yield an improvement over global tying. This is due to the fact that the 5’ bias depends very little on the transcript length, as can be seen in Figure 2 (b), and introducing an additional group of unnecessary parameters into the Mix^2^ model therefore leads to slight overfitting. PennSeq performs better than Cufflinks on data with 3’ bias but as in Figure 3(c) performs worse than Cufflinks in the majority of the cases.

### 3.2 Experiments on the Microarray Quality Control (MAQC) data

This section compares the accuracy of the Mix^2^ model, Cufflinks and PennSeq on two publicly available real RNA-Seq data sets generated from the Universal Human Reference (UHR) RNA and brain (HBR) RNA of the Microarray Quality Control (MAQC) data [10]. These RNA samples were sequenced on an Illumina GenomeAnalyzer resulting in 7 lanes per sample of 35 bps single-end reads [1]. The RNA-Seq data of the MAQC data set were downloaded from the NCBI read archive under accession number SRA010153, while the associated qPCR values were downloaded from the Gene Expression Omnibus under accession number GSE5350. The reads of all 14 lanes were aligned to GRCh37/hg19 with tophat2 [16]. Rather than the RefSeq annotation, Ensembl version 75 was used in the experiments, since the Ensembl annotation contains in many cases more transcripts per gene than RefSeq and therefore yields a more challenging and also larger test set. Since the MAQC data set records the association between qPCR probes and RefSeq annotations it was necessary to select only those qPCR probes whose associated RefSeq annotations have a unique Ensembl equivalent. In addition, genes containing a single transcript were removed from the test set as it is not necessary to estimate relative abundances in this case. This resulted in a test set containing 798 transcripts, more than twice the 331 transcripts of the test set in [4]. On average 8.6 transcripts were contained in each gene of the test set. Since the RNA-Seq data from MAQC are single-end the fragment length *l*(*r*) is unknown and was therefore summed out of (3). This effectively sets the first term of the right-hand side of (3) to 1. As the fragment start the down-stream end of each read was selected. For the Mix^2^ model, the FPKM concentration of transcript *t* = *i* was derived from the relative abundance estimate *α*_*i*_ as follows

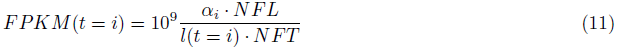

where *NFL* and *NFT* are, respectively, the number of fragments in the locus and the total number of fragments in the experiment. Equation (11) is, strictly speaking, the RPKM [12] and not the FPKM [17] value, as the latter uses a normalized rather than the true transcript length. However, no distinction will be made between the two in the following.

#### 3.2.1 Correlation between FPKM and qPCR values

First, we investigated the correlation between the qPCR values of the MAQC data set and the FPKM values obtained by the Mix^2^ model, PennSeq and Cufflinks. For lane SRR037445 of UHR and lane SRR035678 of HBR this correlation is visualized in Figure 4(a) and Figure 4(b), respectively. The ellipses at the bottom highlight the FPKM values which lie below 10^−3^. Transcripts with such FPKM values were considered not detected and their FPKM values were set to 10^−3^. The average qPCR values of the undetected transcripts are also contained in the graphs. This value should ideally be small since it is more acceptable not to detect transcripts with low rather than high abundance. Figures 4 (a) and (b) show that for Cufflinks the number of points inside the ellipses equals 111 and 127 and thus 14% to 16% of the complete test set are not detected. Similarly, the average qPCR value of between -1.86 and -1.7 is relatively high in comparison to the average for the complete test set, which is -1.02 and -0.99 for the UHR and HBR lane, respectively. In comparison, the number of undetected transcripts is considerably smaller for PennSeq and the Mix^2^ model equaling 17 and 24 or 2% and 3% of the complete test set. In addition, the average qPCR value for the Mix^2^ model is noticeably smaller than for the other methods at -2.55 and -2.83. Figure 4(c) shows that the average R^2^ value is considerably higher for the Mix^2^ model than Cufflinks and PennSeq for both UHR and HBR. In particular, for HBR the R^2^ value for the Mix^2^ model without tying equals 0.54 whereas the R^2^ value for Cufflinks with bias correction and PennSeq is 0.34 and 0.32, respectively. For UHR the R^2^ value for the Mix^2^ model without tying equals 0.60 in comparison to 0.44 for PennSeq and 0.4 for Cufflinks with bias correction, respectively. Figure 4 shows furthermore that group tying gives a small improvement over global tying. However, the Mix^2^ model without any tying performs noticeably better. This indicates that the tying of transcripts which is purely based on gene association and transcript length is too simple for the MAQC data. Figure 4(d) shows also a moderate increase in Spearman correlation by the Mix^2^ model which is more prominent for UHR. For HBR, on the other hand, the difference between 0.69 for Cufflinks with bias correction and 0.71 for the Mix^2^ model is minute. Regarding the accuracy of differential expression calls, the large number of undetected transcripts is an important factor and the R^2^ value is thus a more meaningful measure than the Spearman correlation for the evaluation of correlation.

**Figure 4:**
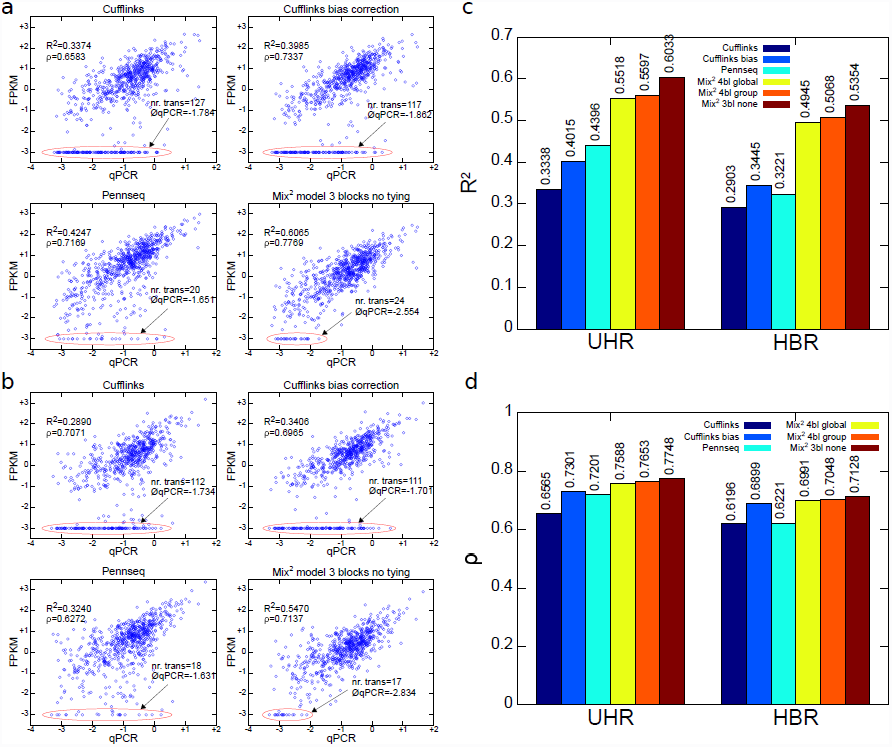
Correlation between qPCR and FPKM values on MAQC data for Cufflinks, PennSeq and Mix^2^ model. Correlation on lane SRR037445 of UHR (a), on lane SRR035678 of HBR (b). Axes are log_10_. FPKM values below 10^−3^ were truncated to 10^−3^ and are highlighted by red ellipses. Number of truncated FPKM values and average of their log_10_(*qPCR*) values at start of arrows. Average of R^2^ value (c) and Spearman correlation *ρ* (d) over all 7 lanes of UHR and HBR.

#### 3.2.2 Correlation between qPCR and FPKM fold change and the detection of differential expression

Fold changes of concentration measures between different samples are used in the analysis of differential expression both for microarrays [18] and RNA-Seq [17]. Therefore, a high correlation between the qPCR and FPKM fold changes is important to ensure accurate differential expression calls. Figure 5(a) shows the correlation between the fold change of the qPCR value on the x-axis and the FPKM value on the y-axis. The fold changes of the FPKM values were calculated between lane SRR037445 of UHR and SRR035678 of HBR. As in the previous section, Figure 5(a) shows for Cufflinks the presence of a large number of transcripts whose FPKM fold change lies considerably above or below the majority of FPKM fold changes. This is visualized by the long straight lines at FPKM fold changes of 10^−4^ and 10^+3^. For PennSeq and the Mix^2^ model, on the other hand, this number is considerably smaller. The R^2^ values and Spearman correlations in Figure 5(a) show that the Mix^2^ model without tying and 3 blocks achieves the best correlation. This is further confirmed by Figure 5(b) which shows the R^2^ value and Spearman correlation averaged over all 49 combinations of the 7 lanes in UHR and HBR. Figure 5(b) shows that the average R^2^ value for the Mix^2^ model is substantially higher at 0.77 than for PennSeq at 0.62 and Cufflinks with bias correction at 0.47. Also the Spearman correlation of the Mix^2^ model at 0.90 is considerably higher than for PennSeq, 0.81, or Cufflinks, 0.76. In order to determine the influence of the correlation of FPKM and qPCR fold changes on the detection of differential expression, a simple classification experiment was performed, similar to an experiment in [1]. Transcripts whose qPCR fold change was above 2 were defined as differentially up-regulated while transcripts whose qPCR fold change was below 0.5 were defined as differentially down-regulated. The remaining transcripts were defined as not differentially expressed. Subsequently all transcripts were classified according to their FPKM fold change. If the FPKM fold change was above a certain threshold the transcript was classified as up-regulated, whereas it was classified as down-regulated if its FPKM fold change was below the inverse of the threshold. The threshold varied between 1.1 and 10^6^ and for each threshold the true and false positive rate, with respect to the qPCR based definitions, were recorded. Here, it was counted as an error if an up-regulated transcript was classified as down-regulated and vice versa. For lane SRR037445 of UHR and SRR035678 of HBR this experiment resulted in the ROC curve in Figure 5(c). This shows clearly that the FPKM values calculated with the Mix^2^ model yield a much better characteristic than the FPKM values calculated with the other methods. Figure 5(d) shows the true positive rate for false positive rates of 0.1, 0.15 and 0.2 averaged over all the 49 combinations of the 7 lanes in UHR and HBR. For a false positive rate of 10%, for instance, the true positive rate for the Mix^2^ model without tying equals 0.71 which is considerably higher than the 0.44 for PennSeq and the 0.24 for Cufflinks. While this difference becomes increasingly smaller for false positive rates of 0.15 and 0.2 it is still prominent with a true positive rate of 0.82 for the Mix^2^ model without tying, 0.68 for PennSeq and 0.65 for Cufflinks for a false positive rate of 0.2.

**Figure 5:**
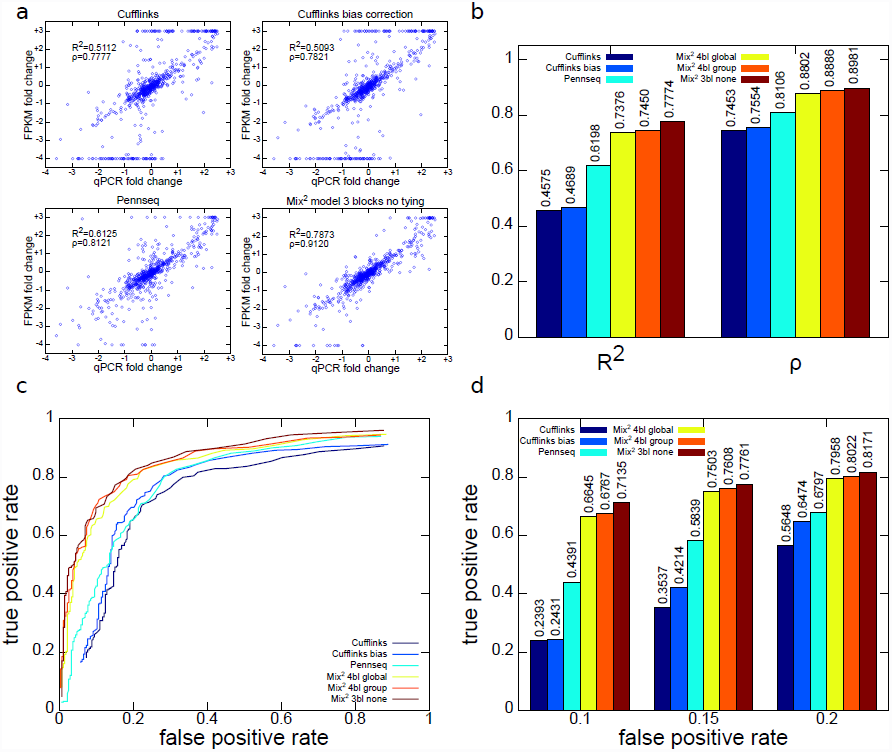
Correlation between qPCR and FPKM fold changes between UHR and HBR for Cufflinks, PennSeq and the Mix^2^ model and ROC curve for classification experiment. (a) Correlation between qPCR and FPKM fold changes between UHR lane SRR037445 and HBR lane SRR035678. (b) Average R^2^ value and Spearman correlation over all 49 combinations of the 7 lanes in UHR and HBR. (c) ROC curve of classification experiment based on FPKM values of UHR lane SRR037445 and HBR lane SRR035678. (d) Average true positive rate over classification experiments for all 49 combinations of lanes in UHR and HBR for false positive rates 0.1, 0.15 and 0.2.

#### 3.2.3 Types of bias in the MAQC data

The Mix^2^ model simultaneously estimates relative abundances and transcript specific fragment distributions and can therefore be used to detect biases present in RNA-Seq data. Figures 6(a) to (f) visualize 6 bias types for a subset of transcripts in one lane of UHR, which were obtained by clustering the fragment start distributions learned by the Mix^2^ model. For this purpose the fragment start distributions of the 798 genes of the MAQC test set from lane SRR037445 of UHR were selected, which were assigned a read count by the Mix^2^ model of at least 100. The minimum read count was set at 100 in order to avoid fragment start distributions which might be inaccurate due to a small amount of data. The 858 fragment start distributions, which satisfied these requirements were then length normalized and hierarchically clustered with UPGMA [15] using the L_1_ distance metric. The resulting tree was traversed from top to bottom and nodes were retained which had at least 5% or 43 of the 858 fragment start distributions. If a node was reduced in size at the next level of the tree without changing the overall shape of the distributions, the top node was chosen. This process resulted in the 6 clusters of fragment start distributions shown on the left side of Figures 6(a) to (f). Figure 6(g), on the other hand, shows the fragment start distribution for the complete unclustered data. The median in Figure 6(g) gives the false impression that the fragment start distributions can be modelled by a uniform distribution. Instead, the distributions separate into a class with 5’ bias, Figures 6(a) to (c), a class with 3’ bias, Figures 6(d) and (e), and a uniform class, Figure 6(f). The classes with 5’ or 3’ bias contain 46.50% of the complete fragment start distributions, while the uniform class contains only 26.92%. Overall, 73.43% of the distributions are contained in one of the classes in Figures 6(a) to (f). The remaining 26.57% of the distributions belong to classes each containing less than 5% of the distributions. Thus, biased fragment distributions represent the majority of the data. There exists, furthermore, no single bias type but multiple biases are observed in the data.

**Figure 6:**
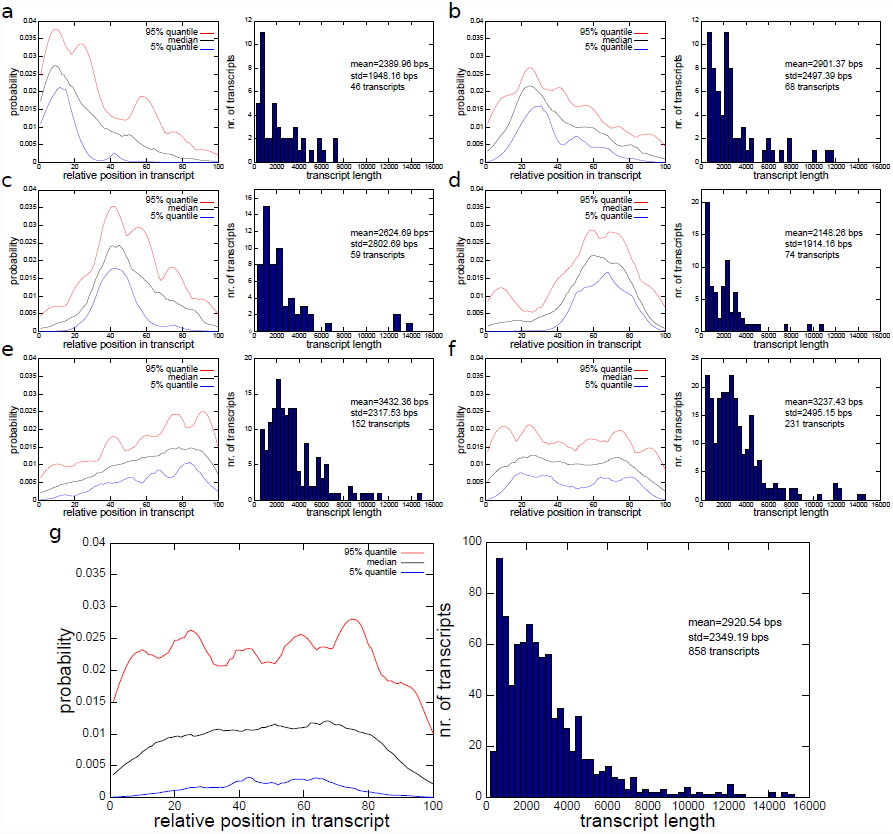
Types of biases detected in lane SRR037445 of UHR and their transcript length distributions. (a) to (f) 6 most prominent biases which account for 73.43% of transcripts. Bias on the left, transcript length distribution on the right. (g) Bias and transcript length distribution of complete, unclustered set of transcripts.

## 4 Discussion

This article introduced the Mix^2^ model which uses a mixture of probability distributions to model the transcript specific fragment distributions in RNA-Seq data. Due to the flexibility of mixture models, the Mix^2^ model can adapt to multiple positional fragment biases of arbitrary complexity. The parameters of the Mix^2^ model are efficiently trained with the EM algorithm yielding simultaneous estimates for the fragment distributions and the relative abundances. In addition, parameters of the Mix^2^ model can be tied between transcripts with similar fragment distribution leading to improved estimates of the relative abundances.

Experiments were conducted on artificial data covering 7 genes of different complexity, 4 types of fragment bias and sample sizes of 500, 1000, 5000 and 10000 fragments. One of the main purposes of these experiments was to optimize the number of mixtures of the Mix^2^ model. These experiments showed that optimization can be performed independent of gene, bias and sample size. A strong dependency on the fragment bias would be problematic as this requires the determination of the bias prior to application of the Mix^2^ model and limits therefore its usefulness as a tool for bias modelling. The optimal number of mixtures for the Mix^2^ model was found to be 3 for a model without tying and any number between 4 and 10 for a Mix^2^ model with tying. These numbers are fairly small given the biases in Figure 2. In particular, it seems rather implausible that a Mix^2^ model with only 3 mixtures should be able to accurately model the 5’+3’ bias in Figure 2(d). This suggests that the potential of the mixtures in the Mix^2^ model has yet to be fully exploited, which might lead to further improvements in the accuracy of the relative abundance estimates. Mix^2^ models with global or group tying appear to be less sensitive to the correct choice of the number of mixtures, whereas Mix^2^ models without tying show a steady decrease in performance for numbers of mixtures beyond the optimum. This might in part be due to the fact that the number of parameters in a Mix^2^ model without tying increases more rapidly with increasing number of mixtures and that it is therefore more prone to overfitting than a Mix^2^ model with parameter tying for a comparable number of mixtures. On the artificial data the Mix^2^ model outperformed both Cufflinks [17] and PennSeq [4] with the exception of data sampled from the Cufflinks bias, for which Cufflinks has the perfect model. This shows that there is still room for improvement for the Mix^2^ model, particularly for larger sample sizes as can be seen from Figure 3(d). The accuracy of PennSeq in the experiments on artificial data was in most cases considerably worse than that of the Mix^2^ model and Cufflinks. This might be due to the uniform sampling with the Dirichlet distribution which results in mainly non-zero abundances. Hence, the overall fragment start distribution of the artificial data in a gene locus can differ strongly from the distribution for individual transcripts. PennSeq, on the other hand, approximates the transcript specific fragment distribution by the part of the overall distribution, which overlaps the transcript, and by the transcript specific reads.

Experiments were also performed on RNA-Seq data generated from the Universal Human Reference (UHR) RNA and brain (HBR) RNA from the Microarray Quality Control (MAQC) data set [10]. These experiments showed considerably better correlation between the qPCR and FPKM values for the Mix^2^ model than for Cufflinks and PennSeq. Furthermore, the correlation between the qPCR and FPKM fold changes between UHR and HBR are noticeably higher for the Mix^2^ model than for the other methods. As shown in a classification experiment, this leads to substantially higher accuracy in the detection of differential expression. In comparison to the experiments on artificial data, on the MAQC data PennSeq performed better than Cufflinks in many cases. This might hint at the fact that, contrary to the sampling for the artificial data, only few abundances in a gene are non-zero and the overall fragment distribution overlapping a transcript is therefore a reasonable approximation for the transcript specific fragment distribution. Overall, the correlation between qPCR and FPKM values found in our experiments agrees reasonably well with the results in [4]. Although in comparison to [4], Cufflinks appears to perform slightly better and PennSeq slightly worse. This discrepancy might be the result of improvements in Cufflinks and the higher complexity of our test set. In contrast to the experiments on artificial data, experiments on the MAQC data showed a degradation in the performance of the Mix^2^ model when tying parameters. This cannot be attributed to a suboptimal choice of parameters in the transcript clustering procedure, which was discussed in Section 2.2. In fact, preliminary experiments showed that changes of the parameters had very little effect on the accuracy of the results. We therefore refrained from optimizing the parameters of the clustering procedure also in light of the large number of experiments needed to cover all parameter combinations in a brute-force approach. Rather than a suboptimal parameter choice, the reason for the degradation in performance seems to be the fact that the similarity of positional fragment bias does not exclusively depend on the gene membership of a transcript and the transcript length. This fact was also highlighted by the experiments in Section 3.2.3. Thus, rather than optimizing the transcript clustering parameters, searching for an extended set of criteria for transcript similarity seems more promising. Further work will be needed to obtain similar improvements through parameter tying on the MAQC data as observed on the artificial data.

Finally, the Mix^2^ model was used to determine the dominant biases in lane SRR037445 of UHR. This was achieved by clustering the transcript specific fragment distributions estimated by the Mix^2^ model. We obtained 6 clusters, containing 73.43% of the distributions, of which 5 clusters, containing 46.50%, exhibited non-uniform distributions. The cluster with uniform distributions contained only 26.92%. Some of these clusters exhibit distinct biases. The presence of a 3’ bias as in Figure 6(e), for instance, is a common feature of library preparations with cDNA fragmentation [20] and the position of the tail-off in the distribution might therefore be related to the distribution of the fragment lengths. The 5’ bias in Figures 6(a) to (c), on the other hand, is potentially the result of RNA degradation. Contrary to Figure 2 there is no obvious relationship between bias and transcript length, although correlations between the two do exist. For instance, transcripts whose fragment start distributions are 3’ biased or uniform as in Figures 6(e) and (f) tend to be longer than the average transcript, whereas transcripts with fragment start distributions in Figures 6(a) and (d) are noticeably shorter. While this is just a first and very cursory glance at the data a more detailed analysis will hopefully reveal relations between biases and RNA sequences of transcripts, which will eventually lead to a better tying strategy for the Mix^2^ model on real RNA-Seq data.

In summary, the Mix^2^ model can be used as an explorative tool to investigate the positional biases present in RNA-Seq data and thereby study the influence of library preparation, sequencing and data processing on the accuracy of transcript concentration estimates. In addition, and more importantly, our results show that in comparison to current state-of-the-art methods the Mix^2^ model yields substantially improved transcript concentration estimates for RNA-Seq data and, as a result, leads to higher accuracy in the detection of differential expression.

## Acknowledgement

Andreas Tuerk was partly funded by the Austrian Research Promotion Agency (FFG) under grant number 838191. Gregor Wiktorin and Serhat Güler were partly funded by the Wiener Arbeitnehmerinnen Förderungsfond (WAFF) under the ’’Förderung Innovation und Beschaftigung” scheme.

We would like to thank Michael Ante for providing valuable input to the manuscript and testing the Mix^2^ model software. Special thanks are due to Lexogen’s founder, Alexander Seitz, for encouraging this project.

## References

[1] James H Bullard, Elizabeth Purdom, Kasper D Hansen, and Sandrine Dudoit. Evaluation of statistical methods for normalization and differential expression in mRNA-Seq experiments. BMC Bioinformatics, 11:94, 2010.

[2] Kasper D Hansen, Steven E Brenner, and Sandrine Dudoit. Biases in Illumina transcriptome sequencing caused by random hexamer priming. Nucleic Acids Res, 38(12):e131, Jul 2010.

[3] L. L. Hsiao, R. V. Jensen, T. Yoshida, K. E. Clark, J. E. Blumenstock, and S. R. Gullans. Correcting for signal saturation errors in the analysis of microarray data. BioTechniques, 32(2), February 2002.

[4] Yu Hu, Yichuan Liu, Xianyun Mao, Cheng Jia, Jane F. Ferguson, Chenyi Xue, Muredach P. Reilly, Hongzhe Li, and Mingyao Li. PennSeq: accurate isoform-specific gene expression quantification in RNA-Seq by modeling non-uniform read distribution. Nucleic Acids Research, 42(3):e20, 2014.

[5] Bo Li and Colin Dewey. RSEM: accurate transcript quantification from RNA-Seq data with or without a reference genome. BMC Bioinformatics, 12(1):323, 2011.

[6] Bo Li, Victor Ruotti, Ron M Stewart, James A Thomson, and Colin N. Dewey. RNA-Seq gene expression estimation with read mapping uncertainty. Bioinformatics, 26(4):493–500, Feb 2010.

[7] Heng Li, Bob Handsaker, Alec Wysoker, Tim Fennell, Jue Ruan, Nils Homer, Gabor Marth, Goncalo Abecasis, Richard Durbin, and 1000 Genome Project Data Processing Subgroup. The Sequence Alignment/Map format and SAMtools. Bioinformatics, 25(16):2078–2079, August 2009.

[8] Jun Li, Hui Jiang, and Wing Wong. Modeling non-uniformity in short-read rates in RNA-Seq data. Genome Biology, 11(5):R50, 2010.

[9] Wei Li and Tao Jiang. Transcriptome assembly and isoform expression level estimation from biased RNA-Seq reads. Bioinformatics, 28(22):2914–2921, 2012.

[10] MAQC Consortium. The MicroArray Quality Control (MAQC) project shows inter-and intraplatform reproducibility of gene expression measurements. Nature Biotechnology, 24(9):1151–1161, September 2006.

[11] Aziz M Mezlini, Eric JM Smith, Marc Fiume, Orion Buske, Gleb Savich, Sohrab Shah, Sam Aparicion, Derek Chiang, Anna Goldenberg, and Michael Brudno. iReckon: Simultaneous isoform discovery and abundance estimation from RNA-seq data. Genome Research, 2012.

[12] Ali Mortazavi, Brian A Williams, Kenneth McCue, Lorian Schaeffer, and Barbara Wold. Mapping and quantifying mammalian transcriptomes by RNA-Seq. Nat Methods, 5(7):621–628, Jul 2008.

[13] Marius Nicolae, Serghei Mangul, Ion Mandoiu, and Alex Zelikovsky. Estimation of alternative splicing isoform frequencies from RNA-Seq data. Algorithms for Molecular Biology, 6(1):9, 2011.

[14] Adam Roberts, Cole Trapnell, Julie Donaghey, John L Rinn, and Lior Pachter. Improving RNA-Seq expression estimates by correcting for fragment bias. Genome Biol, 12(3):R22, Mar 2011.

[15] R. R. Sokal and C. D. Michener. A statistical method for evaluating systematic relationships. University of Kansas Scientific Bulletin, 28:1409–1438, 1958.

[16] Cole Trapnell, Lior Pachter, and Steven L Salzberg. TopHat: discovering splice junctions with RNA-Seq. Bioinformatics, 25(9):1105–1111, May 2009.

[17] Cole Trapnell, Brian A Williams, Geo Pertea, Ali Mortazavi, Gordon Kwan, Marijke J. van Baren, Steven L Salzberg, Barbara J Wold, and Lior Pachter. Transcript assembly and quantification by RNA-Seq reveals unannotated transcripts and isoform switching during cell differentiation. Nat Biotechnol, 28(5):511–515, May 2010.

[18] Virginia Goss Tusher, Robert Tibshirani, and Gilbert Chu. Significance analysis of microarrays applied to the ionizing radiation response. Proceedings of the National Academy of Sciences, 98(9):5116–5121, 2001.

[19] Lin Wan, Xiting Yan, Ting Chen, and Fengzhu Sun. Modeling RNA degradation for RNA-Seq with applications. Biostatistics, 13(4):734–47, 2012.

[20] Zhong Wang, Mark Gerstein, and Michael Snyder. RNA-Seq: a revolutionary tool for transcriptomics. Nat Rev Genet, 10(1):57–63, Jan 2009.

[21] Zhengpeng Wu, Xi Wang, and Xuegong Zhang. Using non-uniform read distribution models to improve isoform expression inference in RNA-Seq. Bioinformatics, 27(4):502–508, Feb 2011.

